# Phosphoproteomic profiling reveals signaling pathways modulated by *Annona muricata* leaf extract in oral adenosquamous carcinoma cells

**DOI:** 10.64898/2026.01.23.701273

**Authors:** Mohd Altaf Najar, Devasahayam Arokia Balaya Rex, Prashant Kumar Modi, G. P Suchitha, S Amrutha, T. S. Keshava Prasad, Shobha Dagamajalu

## Abstract

Phosphorylation driven dysregulation of intracellular signaling networks is a central feature of cancer initiation, progression, and therapeutic resistance. Although *Annona muricata* leaf extracts have demonstrated anticancer activity across multiple experimental models, the underlying molecular mechanisms particularly at the level of phosphorylation dependent signaling remain poorly understood. In this study, we employed a tandem mass tag TMT-based quantitative phosphoproteomic approach to systematically characterize signaling alterations induced by methanolic *Annona muricata*leaf extract (AME) in oral squamous cell carcinoma (OSCC) CAL-27 cells.

Functional assays revealed that AME significantly inhibited cell proliferation, migration, and clonogenic survival. Global phosphoproteomic profiling identified 6,362 phosphopeptides corresponding to 1,964 unique phosphorylation sites across nearly 7,000 phosphoproteins. AME treatment induced widespread, time-dependent hypophosphorylation, indicating a coordinated suppression of oncogenic signaling networks. Pathway and network analyses revealed marked inhibition of signaling pathways associated with key oncogenic kinases, including cyclin-dependent kinases (CDKs), mitogen-activated protein kinases (MAPKs), and signaling modules linked to EGFR and mTOR pathways. Kinase-substrate enrichment and kinome mapping further demonstrated reduced inferred activity of CDK and MAPK driven signaling, accompanied by suppression of cell cycle progression, mitosis, and checkpoint regulation.

Collectively, these findings demonstrate that AME induces systems-level remodeling of phosphorylation dependent signaling networks, enforcing a growth-restrictive cellular state in OSCC cells. This study highlights quantitative phosphoproteomics as a powerful strategy for dissecting natural product mediated regulation of oncogenic signaling and provides mechanistic insight into the anticancer potential of *Annona muricata*.

## Introduction

Cancer remains a leading cause of mortality worldwide [1], with dysregulated cellular signaling representing a central hallmark of tumor initiation and progression [2]. Aberrant phosphorylation of protein kinases and downstream signaling proteins drives uncontrolled proliferation, survival, and metastasis in many cancer types [3]. Consequently, systematic characterization of phosphorylation-dependent signaling networks has become essential for understanding cancer biology [4]. Advances in mass spectrometry-based quantitative phosphoproteomics now enable high-resolution, global interrogation of kinase activity and signaling pathway dynamics in cancer cells.

Despite significant advances in cancer treatment, including surgery, radiotherapy, and chemotherapy, treatment-associated toxicity and therapy resistance remain major challenges. These limitations have spurred interest in biologically active natural products as sources of anticancer agents and molecular probes [5, 6]. *Annona muricata* L., a member of the Annonaceae family, has attracted attention due to reported anti-inflammatory, antiproliferative, and anticancer activities across multiple cancer models [7, 8]. Extracts derived from *A. muricata* leaves have been shown to inhibit cancer cell growth in lung, colon, pancreatic, prostate, and breast cancer systems [8-11].

However, the molecular mechanisms underlying the anticancer effects of *A. muricata* remain incompletely understood. Importantly, concerns regarding neurotoxicity and systemic toxicity associated with prolonged exposure to *A. muricata* derived compounds highlight the need for mechanistic insight rather than empirical application. Understanding how *A. muricata* modulates intracellular signaling pathways is therefore critical for evaluating its biological relevance and potential utility.

In this study, we employed a tandem mass tag TMT-based quantitative phosphoproteomic approach to investigate phosphorylation-driven signaling alterations induced by methanolic *A. muricata* leaf extract (AME) in oral squamous cell carcinoma (OSCC) CAL-27 cells. By integrating phosphoproteomic profiling with functional assays assessing proliferation, migration, and clonogenic potential, we aimed to delineate signaling networks associated with the anticancer effects of AME. This systems-level analysis provides insight into phosphorylation-dependent pathways modulated by *A. muricata* and establishes phosphoproteomics as a powerful strategy for dissecting natural product–mediated signaling in OSCC.

## Materials and Methods

### Reagents

Cell culture-related consumables such as Dulbecco’s modified Eagle medium (DMEM) (Cat#12100046), fetal bovine serum (FBS), and 100× antibiotic/antimycotic solution (Cat#15240062) were procured from Gibco. Phospho-AKT phospho□ERK antibodies were procured from Cell Signaling Technology (Cell Signaling Technology, Beverly, MA). TMT labels and all solvents used in the LC/MS-MS were purchased from Thermo Fisher Scientific.

### Preparation of *A. muricata* methanolic extract (AME)

The leaves of *A. muricata* were obtained from the botanical garden of Yenepoya (Deemed to be University). The identification was performed by Dr. K.R. Chandrashekar, Professor, Department of Applied Botany, Mangalore University, Mangalagangothri - 574 199, Karnataka, India. The plant material was deposited in the Herbarium, Department of Applied Botany, Mangalore University, India. The mature leaves were air-dried for a week and powdered using a grinding mill. Later, powdered samples were extracted with methanol for overnight. The extract was filtered and dried with a speed vac.

### Preliminary screening of phytochemicals

AME were subjected to qualitative phytochemical analysis for the detection of various chemical constituents such as alkaloids, carbohydrates, proteins and amino acids, phenolic contents and tannins according to the method of Raman (2006).

### Determination of total phenolics

Total phenolic compounds were evaluated as per the protocol of Bray and Thorpe [12]. Different concentrations of AME were mixed with Folin & ciocaiteu’s phenol reagent and shaken vigorously. After 2min, 20% Na_2_CO_3_ was added and incubated for 30min at room temperature and absorbance was read at 765nm. Total phenolic content was calculated by using the standard gallic acid.

### Determination of flavonoids

Total flavonoid content was detected by aluminium chloride method [13]. Different concentrations of AME were mixed with 10% aluminium chloride and 1M potassium acetate. The absorbance was measured at 440nm, after incubation for 30min at room temperature. The flavonoid concentration was calculated by using the standard quercetin.

### Determination of total antioxidant capacity (TAC)

Phosphomolybdenum assay was used to measure TAC as per the protocol of Prieto et al [14]. Different concentrations of AME was incubated with 0.5ml of phosphomolybdenum reagent in a water bath at 95°C for 90min. Absorbance was measured at 695nm after it cooled to room temperature. L-Ascorbic acid was used as standard and TAC was indicated as μg ascorbic acid equivalents/g of the extract.

### DPPH free radical scavenging activity

DPPH radical scavenging assay was estimated according to the protocol of Brand-Williams et al., 1995 [15] with slight modification. The different concentrations of AME were mixed with 100µl of 0.1mM DPPH reagent. The absorbance was measured at 517nm after incubating in the dark for 30mins. The inhibition percentage was estimated by:

Inhibition % = (Ac – As)/Ac × 100 where, Ac = Absorbance of the control, As = Absorbance of the test

### Cell culture

The CAL-27 cells were obtained from American Type Culture Collection (ATCC, Manassas, VA) and grown in Dulbecco’s Modified Eagle’s Medium (DMEM) supplemented with 10% fetal bovine serum (FBS) and 1× antibiotic and antimycotic solution. The cells were cultured in a humidified incubator at 37°C in an atmosphere of 5% CO_2_. Cells were grown up to 70-90% confluence and used for MTT assay.

### MTT cell proliferation assay

To study the effect of AME on cell proliferation of CAL-27 cells, an MTT assay was carried out as described in the literature [16]. CAL-27 cells were seeded at a density of 1 x 10^4^ cells/well in a 96-well plate and treated with AME at a varying concentration from 0-2000µg/ml for 48hr. After incubation, MTT was added and incubated for 4hr at 37^0^C. The reduced formazan product was dissolved in DMSO: absolute ethanol (1:1) and dye intensity was read at 570 and 650nm using a multimode plate reader (Multiskan sky, Thermo Fisher Scientific). Each treatment was carried out in triplicates.

### Colony formation assay

A colony formation assay was performed to assess the colony-forming ability of cells as discussed previously [17]. The cells were seeded at a density of 3 × 10^3^ cells/well in a six-well plate. After 24hr, cells were treated with AME, and cultured for 10 days with changing the media. The cell colonies were fixed and visualized by crystal violet staining (Sigma□Aldrich). The extra dye was removed and colonies were counted. All experiments were done in three biological replicates. Images were taken with an inverted light microscope (Carl Zeiss, USA) at 8× magnification.

### Wound healing scratch assay

Wound healing scratch assay was carried out as described in the literature [18]. Approximately 3 × 10^5^ cells/well were cultured in a six-well plate and incubated until it becomes 80% confluence. The uniform wound was then introduced and incubated in the presence and absence of AME. The cells that migrated into the wound region were determined and images were captured under a phase-contrast microscope.

### Western blot analysis

To check the inhibitory time points of AME on CAL-27 cells, the cells were treated with AME when it reached 70% confluence and harvested at different time intervals of 5min, 15min, 30min, 1hr, 2hr and 3hr. The cells were washed with chilled 1X PBS and lysed using cell lysis buffer [50mM triethyl ammonium benzyl chloride (TEABC), 2% SDS, 1mM Sodium orthovanadate, 2.5mM Sodium pyrophosphate, 1mM β-Glycero phosphate]. The cell lysates were probe sonicated on ice using Q□Sonica (Cole□Parmer). The supernatant was used to quantify the protein by using bicinchoninic acid (BCA) protein estimation kit. AKT and MAPK proteins were assessed. Western blotting was carried out as described previously ((Modi et al., 2012). β-actin was used as a loading control.

### Sample preparation for LC-MS/MS phosphoproteomic analysis

The CAL-27 cells were grown until 70% confluence and treated with AME in different time points for 5min, 15min, 30min, 1hr and 2hr. The cells were washed with cold 1X PBS thrice and harvested with fresh lysis buffer

### Cell lysis and protein digestion

Protein extraction was carried out as described previously [19]. Protein concentration was estimated by using a BCA assay kit. An equal amount of protein from each sample was taken and subjected to reduction with 10mM dithiothreitol (DTT) and alkylation with 20mM iodoacetamide (IAA). Acetone precipitation was performed to remove SDS and the other salts present in the solution. Trypsin digestion was carried out using modified sequence-grade trypsin (Promega, Madison, WI). The clean peptides were dried and stored at −20°C until TMT labeling.

### Tandem mass tag (TMT) labelling and basic pH reversed-phase liquid chromatography (bRPLC)

The resulting peptides were labelled using TMT 10 plex reagents (Thermo Fisher Scientific) as described previously (Karthikkeyan et al., 2020). The peptides were labeled with TMT tags, control with 127N, 5min with 127C, 15min with 128N, 30min with 129N, 1hr with 130N and 2hr with 131. The labeled peptides were equally pooled and vaccum dried before subject into fractionation. The pooled peptides were solubilized in solvent A (10 mM TEABC in water; pH 8.5) and separated on Xbridge BEH C18 Column (Waters Corporation, Milford, MA, USA) in a 1290 Infinity HPLC system (Agilent, Santa Clara, CA, USA). Fractionation was performed as described previously [20]. The samples were concatenated to 12 fractions and vacuum-dried before subjected into phosphopeptide enrichment.

### Phosphopeptide enrichment by refined 2,5-dihydroxybenzoic acid (DHB) method and LC-MS/MS analysis

Phosphopeptide enrichment was carried out by TiO2 DHB strategy as described previously [20, 21]. The enriched peptides were vaccum dried and reconstituted in 0.1% formic acid before inject into LC-MS/MS. Thermo Scientific Orbitrap Fusion Tribrid mass spectrometry (Thermo Fisher Scientific, Bremen, Germany) connected to Easy-nLC-1200 nanoflow liquid chromatography system (Thermo Scientific, Bremen, Germany) was used for data acquisition. The LC-MS/MS data was acquired as described previously [22].

### Data analysis

LC-MS/MS raw data were searched using the software Proteome Discoverer version 2.2 (Thermo Scientific, Bremen, Germany). The MS/MS data were searched against the human protein database (Refseq. 102 databases along with known mass spectrometry contaminants) using search algorithms SEQUEST and Mascot. Search parameters including static and dynamic modifications, minimum peptide length, mass tolerance and false discovery rate was set as described previously [20]. Trypsin was specified as the protease. Perseus 80 tool was used for computing the fold change (FC), the p-value, and the q value. The Sankey diagram was generated using an online Sankey generator (https://sankey□diagram□generator.acquireprocure.com/). Proteins were classified and categorized based on Gene Ontology analysis using DAVID 8.6 functional annotation tool (https://david.ncifcrf.gov/summary.jsp) and pathway analysis using Reactome 83 (https://reactome.org/) tools.. The KinMap tool (http://www.kinhub.org/kinmap/index.html) was used to build the kinome map. The list of identified kinases was searched and differentially regulated kinases were highlighted on the kinome map. Kinase-substrate enrichment analysis (KSEA) was done using the online eXpression2Kinases (X2K; http://amp.pharm.mssm.edu/X2K/) tool.22.

## Results

The literature suggests that *Annona muricata* leaves possess potent anticancer activity, with multiple studies reporting cytotoxic effects across diverse cancer types [23, 24]. However, while these studies demonstrate growth inhibition and induction of cell death, a comprehensive understanding of the underlying signaling mechanisms and molecular targets particularly at the phosphorylation level remains limited.

To elucidate the mechanism of action of *A. muricata* extract in oral squamous cell carcinoma (OSCC), CAL-27 cells were treated with *A. muricata* methanolic extract (AME). Anticancer bioassays were performed to assess its effects on cellular proliferation, migration, and clonogenic potential, followed by a TMT-based quantitative phosphoproteomic analysis to systematically characterize AME-induced alterations in phosphorylation-dependent signaling networks.

### Phytochemical screening of *A. muricata* extract

Preliminary qualitative phytochemical screening of AME revealed the presence of alkaloids, carbohydrates, proteins, and phenolic compounds (Table 1). Quantitative analysis showed that total phenolic and flavonoid contents increased in a concentration-dependent manner (Fig. 1A, B). AME exhibited a progressive increase in total antioxidant capacity, reaching a maximum value of 62.07 ± 1.41 at 800 μg/ml (Fig. 1C). Consistently, DPPH radical scavenging activity increased significantly at higher concentrations, with 65.23 ± 1.07% and 72.24 ± 1.10% scavenging observed at 600 and 800 μg/ml, respectively (Fig. 1D).

**Table 1.**
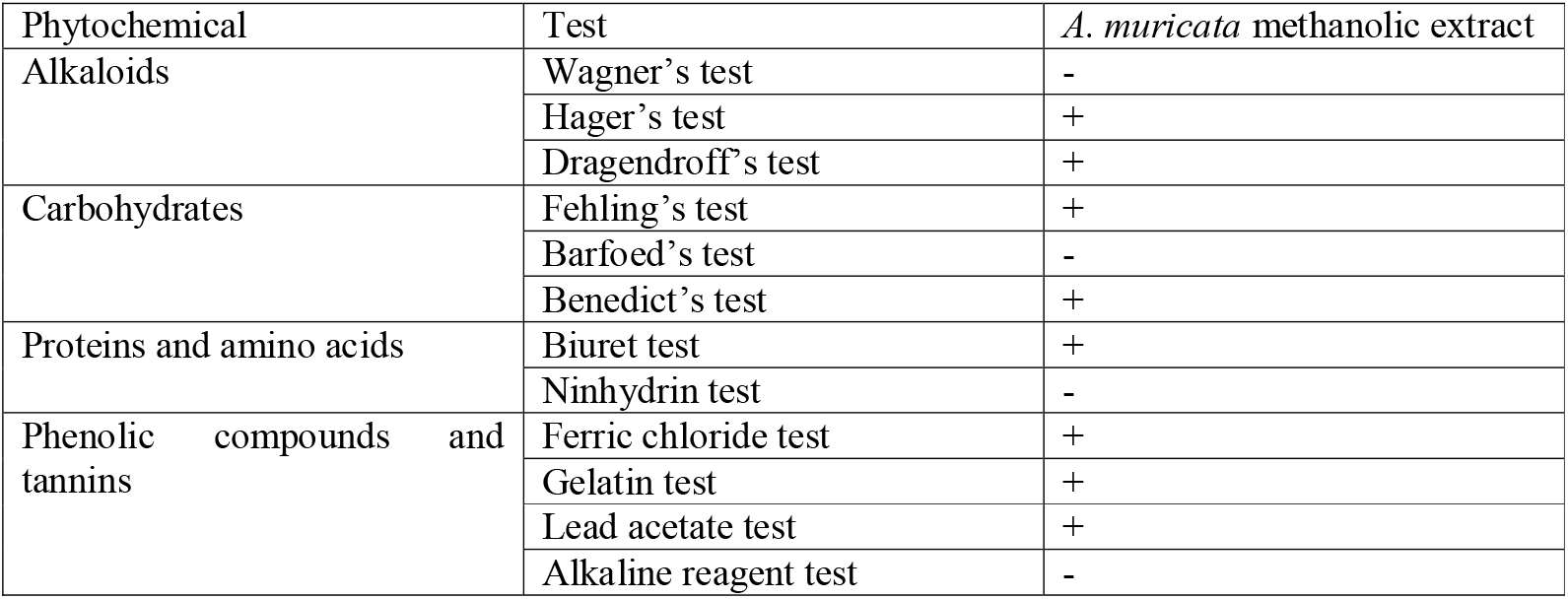
Qualitative phytochemical analysis of the *A. muricata* methanolic extract. Extract showed the presence of alkaloids, carbohydrates, proteins and phenolics as the major constituents.

| Phytochemical | Test | *A. muricata* methanolic extract |
| --- | --- | --- |
| Alkaloids | Wagner's test | - |
|  | Hager's test | + |
|  | Dragendroff's test | + |
| Carbohydrates | Fehling's test | + |
|  | Barfoed's test | - |
|  | Benedict's test | + |
| Proteins and amino acids | Biuret test | + |
|  | Ninhydrin test | - |
| Phenolic compounds and tannins | Ferric chloride test | + |
|  | Gelatin test | + |
|  | Lead acetate test | + |
|  | Alkaline reagent test | - |

**Figure 1.**
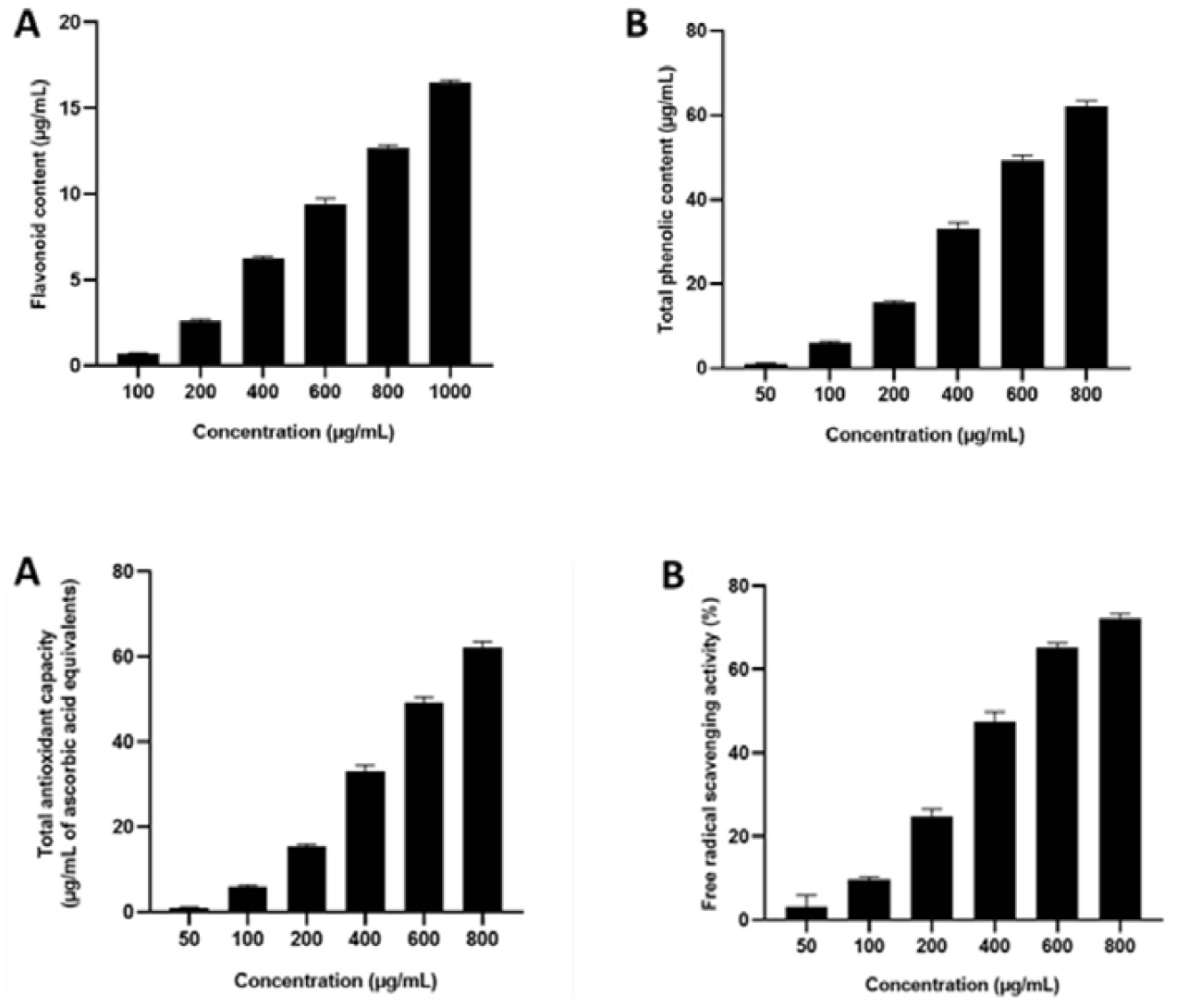
Phytochemical composition and antioxidant activity of *Annona muricata* methanolic extract (AME). (A) Total phenolic content of AME measured at increasing concentrations, showing a concentration-dependent increase. (B) Total flavonoid content of AME at different concentrations. (C) Total antioxidant capacity of AME expressed as antioxidant activity units, demonstrating a progressive increase with extract concentration. (D) DPPH radical scavenging activity of AME at indicated concentrations, showing enhanced free radical scavenging at higher doses. Data are presented as mean ± SEM of three independent experiments.

### AME suppresses growth and survival-associated phenotypes in CAL-27 cells

The anti-proliferative effects of AME on CAL-27 cells were evaluated using the MTT assay. AME treatment resulted in a dose-dependent reduction in cell viability over the concentration range of 500–2000 µg/ml, with an IC□□ value of approximately 900 µg/ml (Fig. 2A). Consistent with these findings, colony formation assays demonstrated a marked reduction in clonogenic survival of AME-treated cells compared to untreated controls (Fig. 2B).

**Figure 2.**
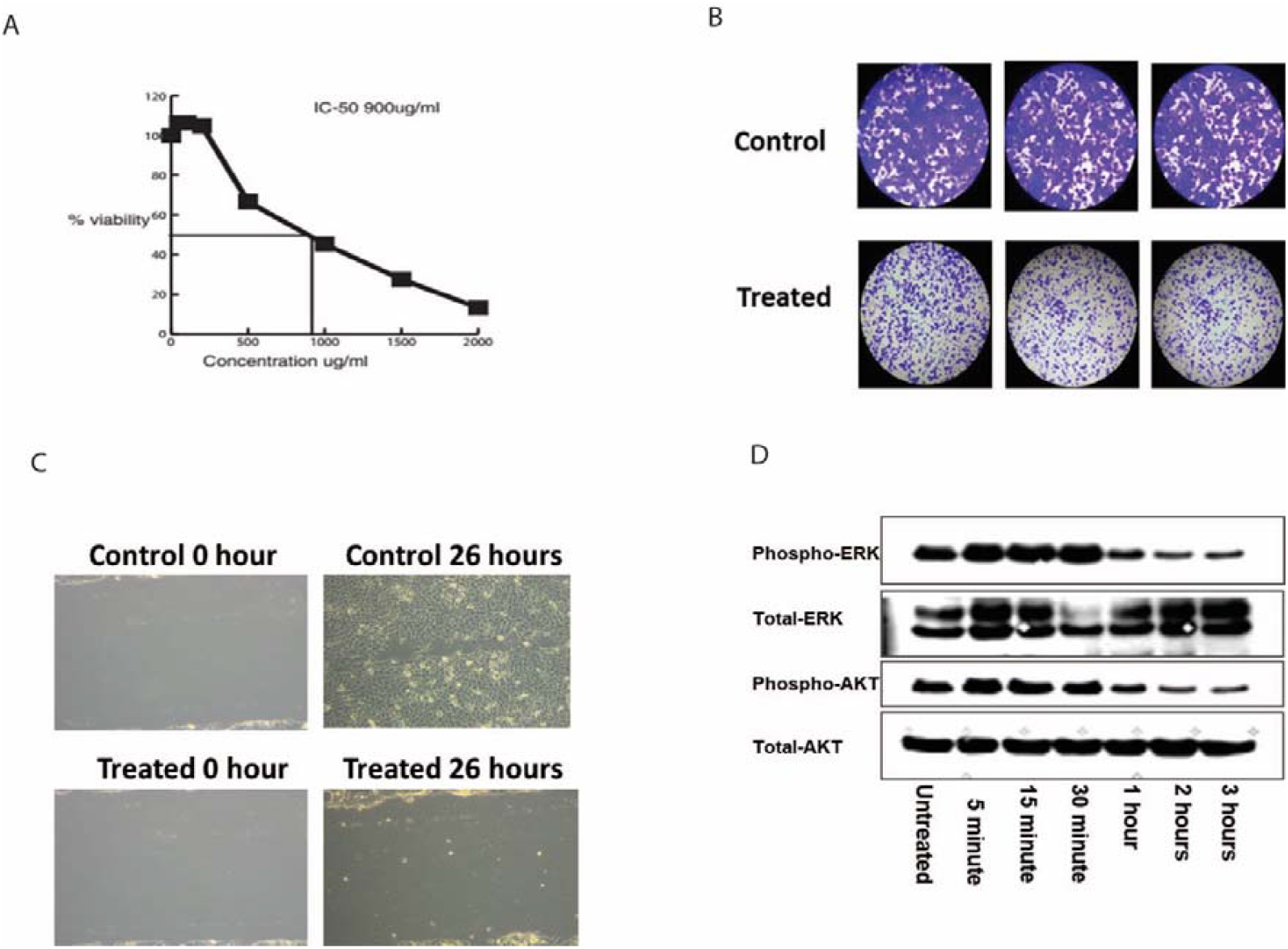
AME suppresses growth, clonogenic survival, migration, and oncogenic signaling in CAL-27 cells. (A) Dose-dependent inhibition of CAL-27 cell viability following treatment with *Annona muricata* methanolic extract (AME) for 24 h, as assessed by MTT assay. The IC□□ value was approximately 900 µg/ml. (B) Representative images and quantification of colony formation assay showing reduced clonogenic survival in AME-treated CAL-27 cells compared to untreated controls. (C) Wound-healing assay demonstrating impaired migratory capacity of CAL-27 cells following AME treatment. Representative images at 0 h and 26 h are shown. (D) Immunoblot analysis showing reduced phosphorylation of AKT and ERK following 1 h of AME treatment, while total AKT and ERK protein levels remained unchanged. Data are presented as mean ± SEM of three independent experiments. Statistical significance was determined using appropriate tests as described in the Methods section.

The effects of AME on cell migration were assessed using a wound-healing assay. Untreated CAL-27 cells exhibited near-complete wound closure within 26 h, whereas AME-treated cells showed a pronounced inhibition of migratory capacity (Fig. 2C). Collectively, these results indicate that AME suppresses cell viability, proliferation, migratory potential, and colony-forming ability of CAL-27 cells in vitro.

To explore potential signaling alterations associated with AME treatment, phosphorylation levels of AKT and ERK were examined. A reduction in phosphorylated AKT and ERK was observed following 1 h of AME exposure, while total protein levels remained unchanged (Fig. 2D), suggesting an early modulation of survival and proliferation-associated signaling pathways.

### Quantitative phosphoproteomic analysis of OSCC cells following A. muricata treatment

To systematically characterize phosphorylation-dependent signaling alterations induced by *A. muricata* extract (AME), we performed a quantitative phosphoproteomic analysis in CAL-27 cells following short-term treatment. A schematic overview of the phosphoproteomic workflow is shown in Figure 3A. In total, we identified 6,362 phosphopeptides corresponding to 1,964 phosphoproteins at a peptide false discovery rate (FDR) of <1% (Supplementary Table 1). These phosphopeptides mapped to 7,002 unique phosphosites. Notably, short-term AME treatment did not result in appreciable changes at the total proteome level, indicating that the observed effects predominantly reflect phosphorylation dynamics rather than changes in protein abundance.

**Figure 3.**
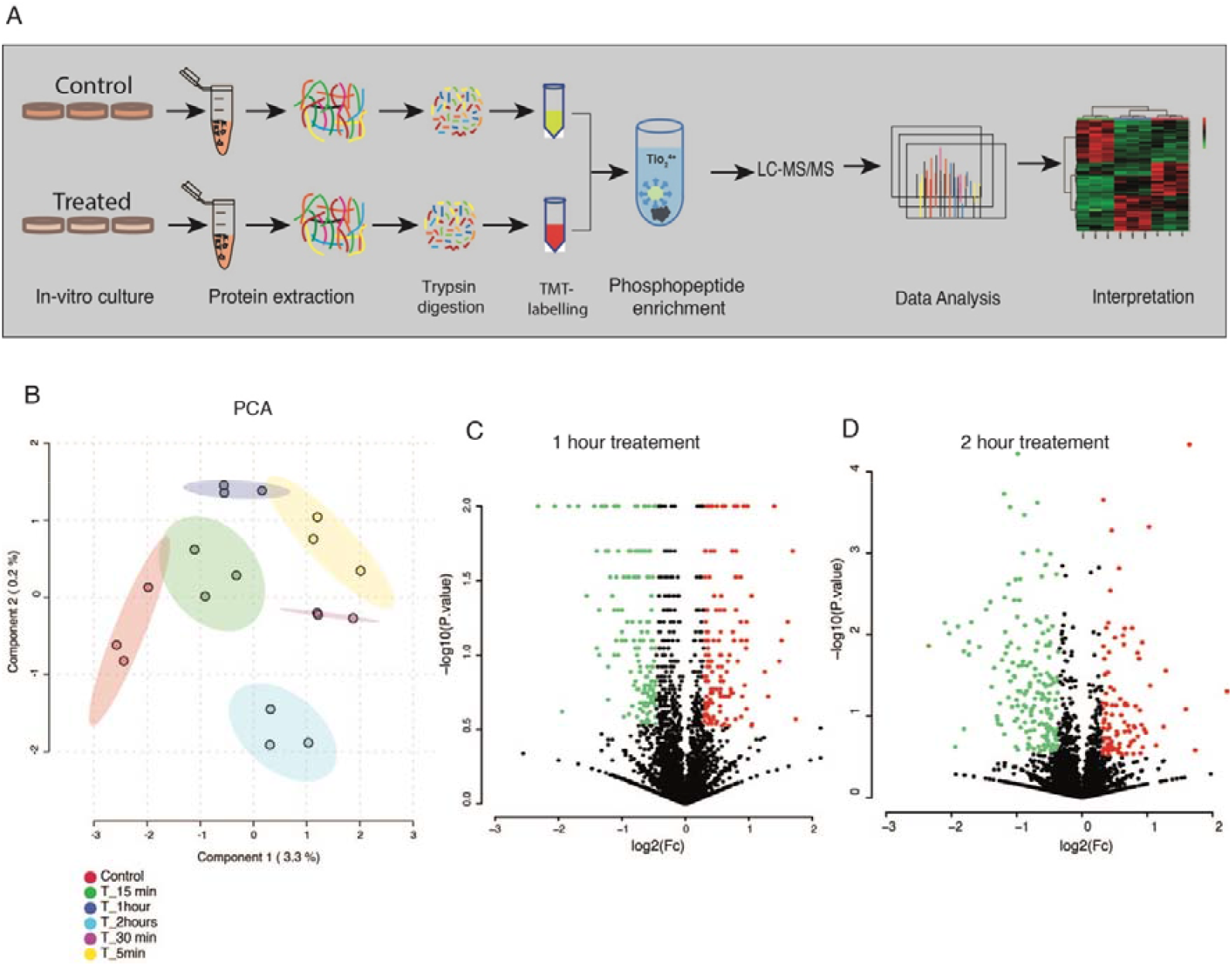
Quantitative phosphoproteomic profiling reveals time-dependent phosphorylation dynamics following AME treatment in CAL-27 cells. (A) Schematic overview of the quantitative phosphoproteomic workflow used to characterize phosphorylation-dependent signaling alterations induced by *Annona muricata* methanolic extract (AME) in CAL-27 cells. (B) Principal component analysis (PCA) of phosphoproteomic datasets showing distinct clustering of samples according to AME treatment time points, indicating reproducible and time-dependent phosphorylation changes. (C, D) Volcano plots depicting differentially regulated phosphopeptides at 1 h (C) and 2 h (D) following AME treatment. Phosphopeptides with increased phosphorylation (fold change ≥1.5) and decreased phosphorylation (fold change ≤0.66) are highlighted. Phosphopeptides were identified at a peptide false discovery rate (FDR) of <1%. Statistical thresholds and filtering criteria are described in the Methods section.

Differential phosphorylation analysis was performed using a fold-change threshold of ≥1.5 for increased phosphorylation and ≤0.66 for decreased phosphorylation. This analysis revealed widespread hypophosphorylation following AME treatment, with 151, 184, 221, 333, and 314 phosphoproteins showing reduced phosphorylation at 5 min, 15 min, 30 min, 1 h, and 2 h, respectively. In contrast, hyperphosphorylation was observed in 65, 37, 88, 120, and 112 phosphoproteins at the corresponding time points. A comprehensive list of differentially regulated phosphopeptides is provided in Supplementary Table 2.

Principal component analysis demonstrated distinct clustering of samples across treatment time points, indicating reproducible and time-dependent phosphoproteomic changes upon AME exposure (Figure 3B). Volcano plot analysis highlighted pronounced phosphorylation changes at 1 h and 2 h of treatment (Figure 3C, 3D). Consistently, hierarchical clustering revealed that the majority of differentially regulated phosphoproteins were observed at these later time points (Figure 4A). Comparative analysis between the 1 h and 2 h treatment conditions identified 146 commonly hypophosphorylated and 46 commonly hyperphosphorylated phosphoproteins (Figure 4B).

**Figure 4.**
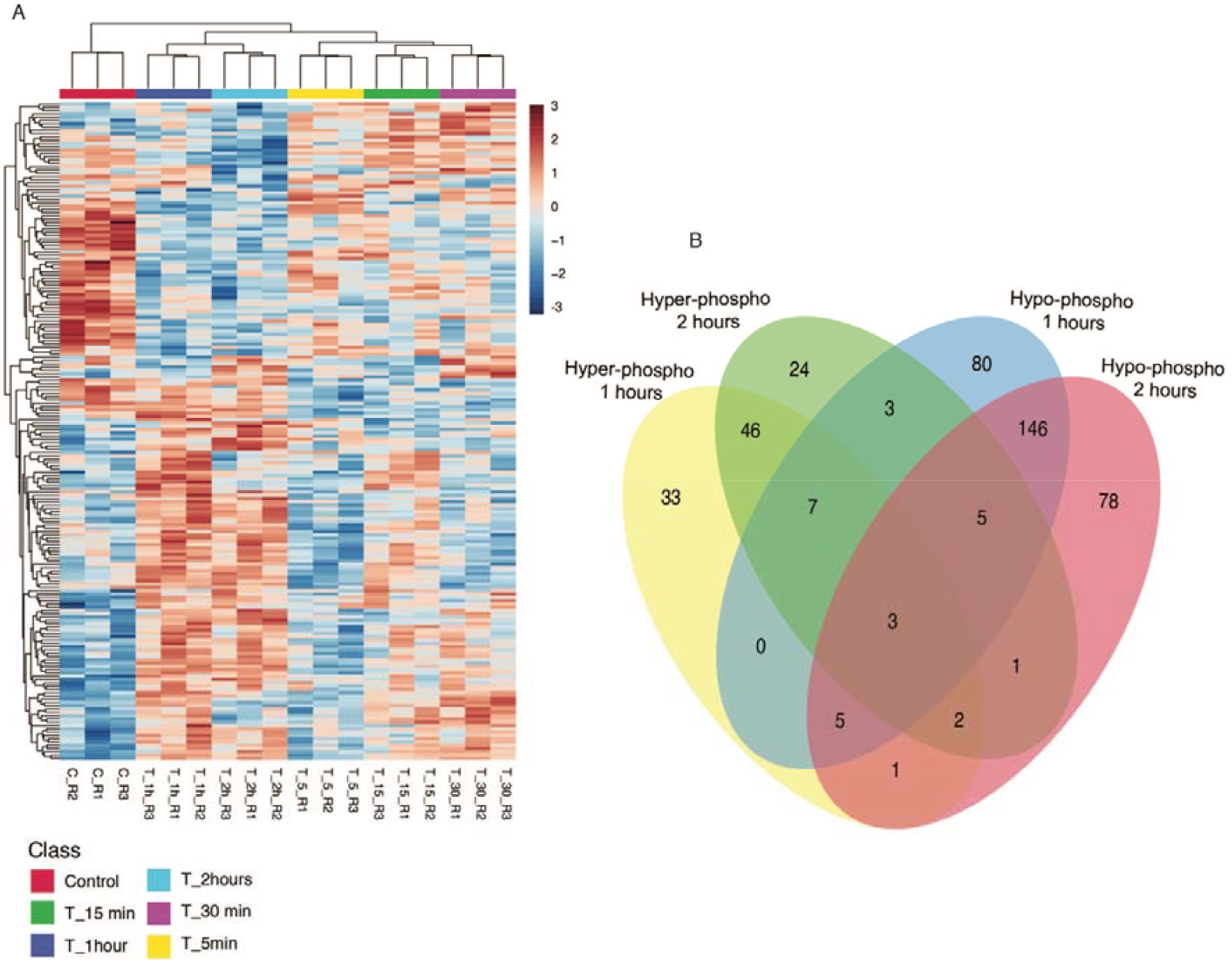
Global landscape of differentially regulated phosphoproteins following short-term AME treatment. (A) Hierarchical clustering heatmap of differentially regulated phosphoproteins across all AME treatment time points (5 min, 15 min, 30 min, 1 h, and 2 h), demonstrating a predominance of hypophosphorylation at later time points. (B) Venn diagram illustrating the overlap of differentially regulated phosphoproteins between 1 h and 2 h AME treatment conditions, highlighting 146 commonly hypophosphorylated and 46 commonly hyperphosphorylated phosphoproteins. Differential phosphorylation was defined using fold-change thresholds of ≥1.5 for increased phosphorylation and ≤0.66 for decreased phosphorylation.

### Functional enrichment analysis of the AME-regulated phosphoproteome

Functional enrichment analysis was performed to characterize the biological relevance of AME-regulated phosphoproteins. Gene Ontology (GO) cellular component analysis indicated that both hypo and hyperphosphorylated proteins were predominantly localized to nuclear and cytoplasmic compartments, including the nucleus, nucleoplasm, cytoplasm, and cytosol. GO biological process analysis revealed that hypophosphorylated proteins were primarily associated with cell-cell adhesion and regulation of transcription, whereas hyperphosphorylated proteins were also enriched for cell-cell adhesion related processes. At the molecular function level, both hypo and hyperphosphorylated proteins were mainly annotated with protein-binding activity (Supplementary Fig. 1).

Pathway enrichment analysis using the Reactome database revealed that hypophosphorylated proteins were significantly enriched in cell-cycle–related pathways, including mitotic progression, nuclear envelope breakdown, and checkpoint regulation, along with signaling pathways involving Rho GTPases, Miro GTPases, and RHOBTB3. Additional enrichment was observed for pathways associated with cellular senescence, oxidative stress-induced senescence, SUMOylation by E3 ligases, and formation of the cornified envelope (Fig. 5A). In contrast, hyperphosphorylated proteins were predominantly enriched in pathways related to RNA metabolism, including mRNA processing and deadenylation, as well as signaling pathways involving Rho GTPases, mTOR and mTORC1 signaling, mitotic spindle checkpoint control, DNA damage- and telomere stress-induced senescence, and ERBB family-mediated signaling (Fig. 5B).

**Figure 5.**
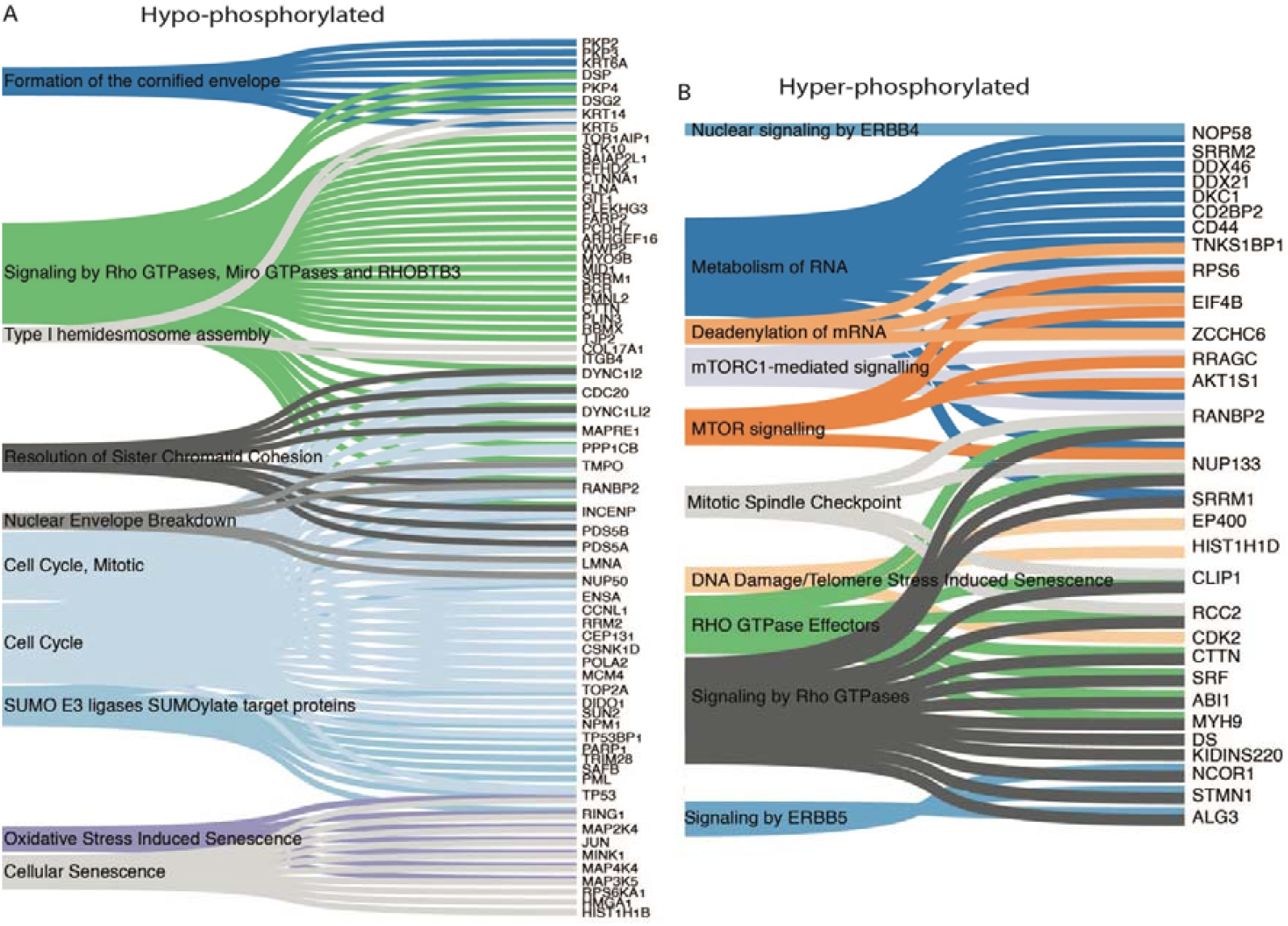
Pathway enrichment analysis of AME-regulated phosphoproteins. (A) Reactome pathway enrichment analysis of hypophosphorylated proteins following AME treatment, showing significant enrichment of cell cycle–related pathways, including mitotic progression, nuclear envelope breakdown, and checkpoint regulation, as well as signaling pathways involving Rho GTPases, Miro GTPases, RHOBTB3, cellular senescence, oxidative stress–induced senescence, SUMOylation by E3 ligases, and formation of the cornified envelope. (B) Reactome pathway enrichment analysis of hyperphosphorylated proteins, revealing enrichment of pathways associated with RNA metabolism (including mRNA processing and deadenylation), mTOR and mTORC1 signaling, mitotic spindle checkpoint control, DNA damage- and telomere stress–induced senescence, Rho GTPase signaling, and ERBB family–mediated signaling. Pathway enrichment was performed using the Reactome database, and statistically significant pathways are shown based on adjusted p-values as described in the Methods section.

Protein-protein interaction network analysis further revealed that hypophosphorylated proteins formed densely connected interaction networks enriched for regulators of the cell cycle and cancer-associated processes (Fig. 6A). Conversely, hyperphosphorylated proteins clustered into interaction networks associated with RNA processing and apoptotic signaling pathways (Fig. 6B).

**Figure 6.**
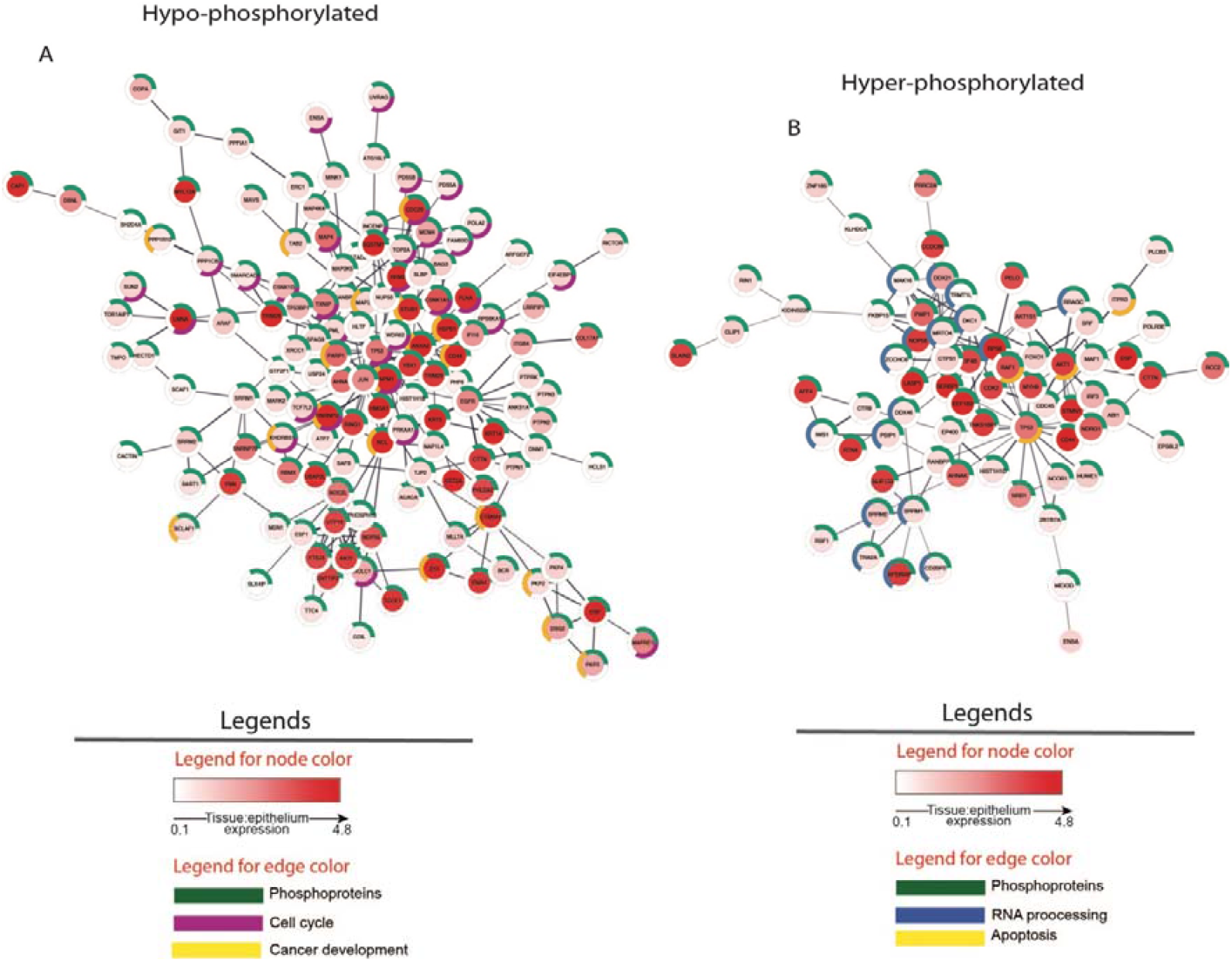
Protein–protein interaction network analysis of AME-regulated phosphoproteins. (A) Protein– protein interaction (PPI) network of hypophosphorylated proteins following AME treatment, revealing densely connected interaction modules enriched for regulators of cell cycle progression and cancer-associated signaling pathways. (B) PPI network of hyperphosphorylated proteins, showing interaction clusters predominantly associated with RNA processing, mRNA metabolism, and apoptotic signaling pathways. Interaction networks were generated using established PPI databases and visualized as described in the Methods section.

### AME alters oncogenic kinase and phosphatase signaling networks in OSC

To investigate the impact of AME on kinase and phosphatase signaling networks, we systematically analyzed phosphorylation changes in kinases and kinase-associated proteins across multiple time points. A total of 30, 32, 38, 64, and 51 kinases or kinase-related proteins exhibited hypophosphorylation at 5 min, 15 min, 30 min, 1 h, and 2 h of treatment, respectively, whereas 15, 3, 13, 23, and 23 showed hyperphosphorylation at the corresponding time points. Several kinase-related proteins, including BAIAP2L1, MARCKS, GIGYF2, EGFR, ARAF, ARHGEF16, MAPRE1, PKP2, CDK12, and NUCKS1, were consistently hypophosphorylated across all treatment durations, while EPS8L2 and CDK16 were consistently hyperphosphorylated.

Analysis of phosphatases revealed hypophosphorylation of GTF2F1 and PPFIA1, whereas ENSA exhibited increased phosphorylation following AME treatment. All hypo and hyperphosphorylated kinases were mapped onto the human kinome tree to visualize global kinase signaling alterations (Fig. 7).

**Figure 7.**
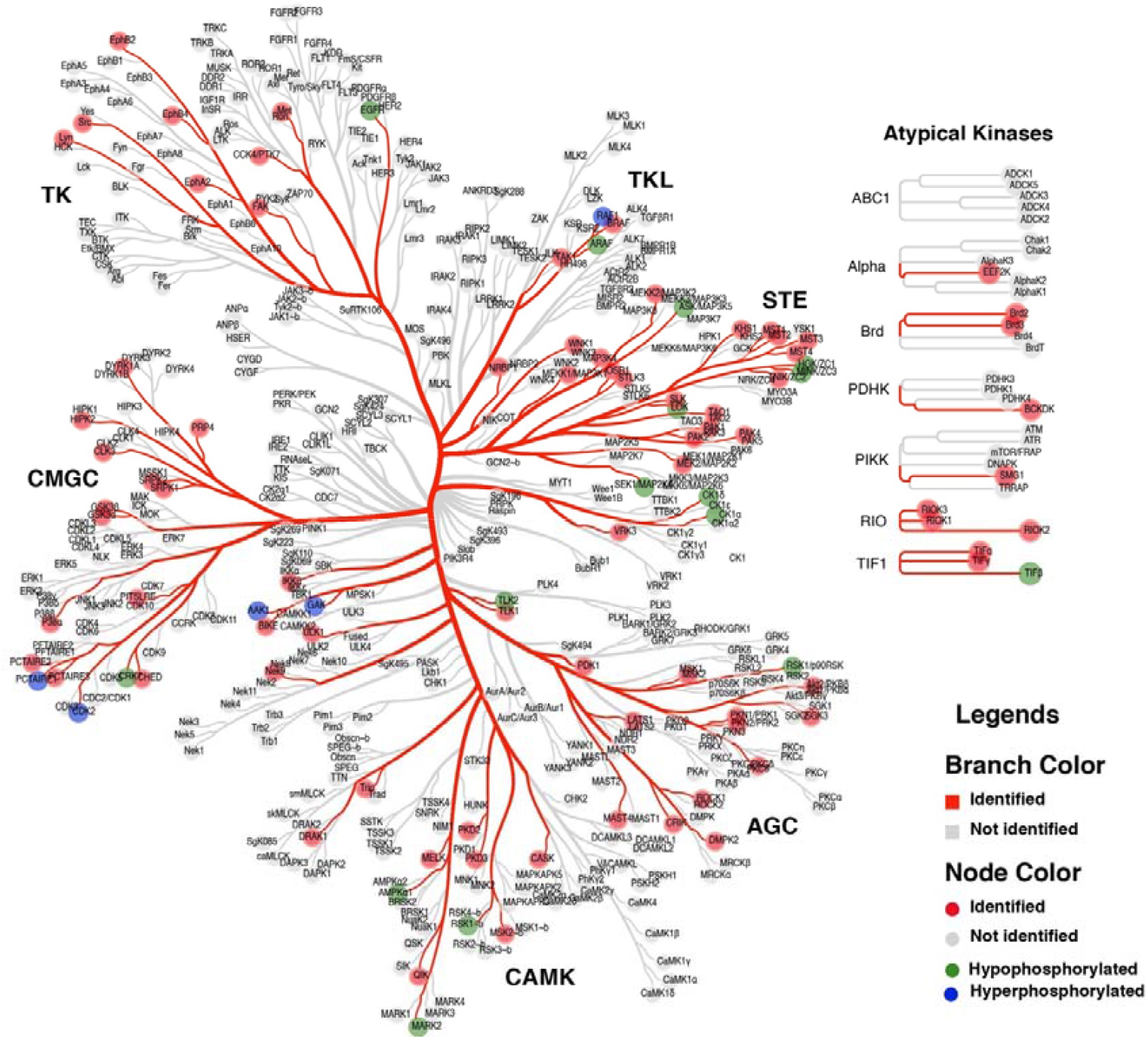
Global kinome landscape of AME-regulated kinase phosphorylation. Kinases and kinase-associated proteins exhibiting differential phosphorylation following AME treatment were mapped onto the human kinome tree. Hypophosphorylated kinases are indicated in blue, while hyperphosphorylated kinases are shown in red. The analysis reveals widespread modulation of kinase signaling networks across multiple kinase families following AME exposure, highlighting a predominance of hypophosphorylation among growth- and cell cycle–associated kinases. Kinase families are organized according to standard kinome classification.

To infer changes in kinase signaling activity, kinase–substrate enrichment analysis (KSEA) was performed. This analysis revealed reduced inferred activity of cyclin-dependent kinases (CDK1, CDK2, and CDK5), as well as mitogen-activated protein kinases including MAPK1, MAPK10, MAPK14, MAP2K1, and MAP2K6. Additional kinases exhibiting reduced inferred activity included PRKCD, TTK, and CSNK1A1. In contrast, increased phosphorylation of substrates associated with kinases involved in survival and cytoskeletal regulation, including PDK1, ROCK1, AKT1/2, mTOR, PKC family members, PAK1/2, RPS6KA2, RPS6KB1, CSNK2A1, and CLK1, was observed (Fig. 8A).

**Figure 8.**
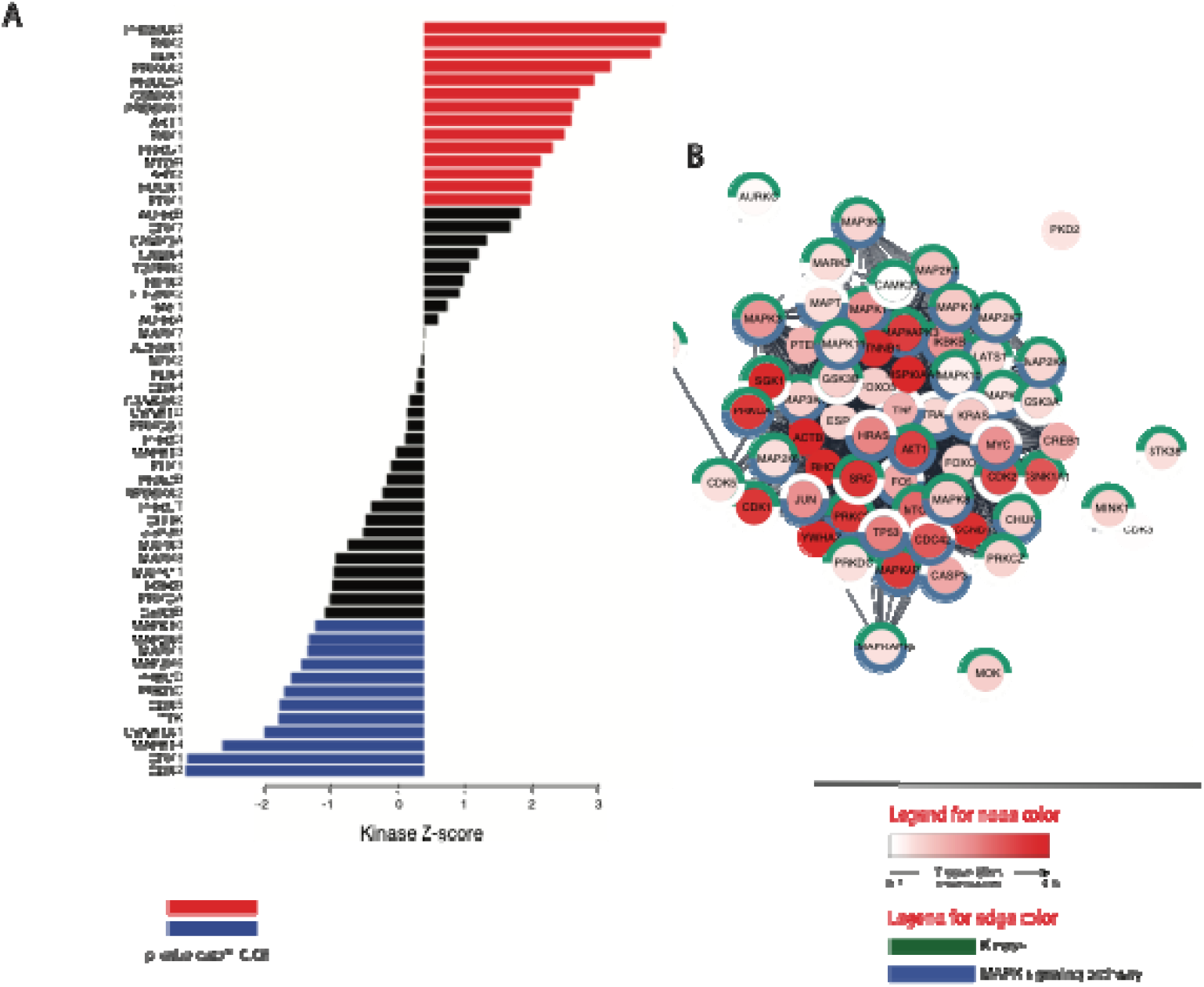
Kinase activity inference and interaction network analysis of AME-regulated kinase signaling. (A) Kinase–substrate enrichment analysis (KSEA) showing inferred changes in kinase activity following AME treatment. Reduced inferred activity was observed for cyclin-dependent kinases (CDK1, CDK2, and CDK5), mitogen-activated protein kinases (MAPK1, MAPK10, MAPK14, MAP2K1, and MAP2K6), and additional kinases including PRKCD, TTK, and CSNK1A1. Increased inferred activity was detected for kinases associated with survival and cytoskeletal regulation, including PDK1, ROCK1, AKT1/2, mTOR, PKC family members, PAK1/2, RPS6KA2, RPS6KB1, CSNK2A1, and CLK1. (B) Protein–protein interaction (PPI) network of hypophosphorylated kinases and kinase-associated proteins, revealing densely connected interaction modules enriched for components of the MAPK signaling pathway, consistent with suppression of oncogenic growth signaling following AME treatment. Kinase activity inference and network construction were performed as described in the Methods section.

Protein-protein interaction network analysis further demonstrated that hypophosphorylated kinases and kinase-associated proteins formed densely connected networks enriched for components of the MAPK signaling pathway, consistent with suppression of oncogenic growth signaling following AME treatment (Fig. 8B).

### AME induces coordinated hypophosphorylation of cell cycle and checkpoint signaling networks in OSCC

To further delineate signaling pathways selectively suppressed by AME, we performed integrative enrichment analyses using ShinyGO 0.85.1 online tool (https://bioinformatics.sdstate.edu/go/) with Kinase library 2023 and KEA 2015 as databases and focusing exclusively on hypophosphorylated phosphoproteins. Kinase enrichment analysis revealed significant overrepresentation of substrates associated with multiple cyclin-dependent kinases, including CDK1, CDK2, CDK7, CDK11A, CDK15, and CDK18, along with key mitotic and checkpoint regulators such as AURKB, CHEK1, CHEK2, and PRKDC. In addition, hypophosphorylated substrates corresponding to mitogen-activated protein kinases (MAPK1, MAPK3, MAPK8, MAPK9, and MAPK14) and mTOR were prominently enriched, indicating broad attenuation of proliferative and survival-associated kinase signaling (Fig. 9A &B).

**Figure 9.**
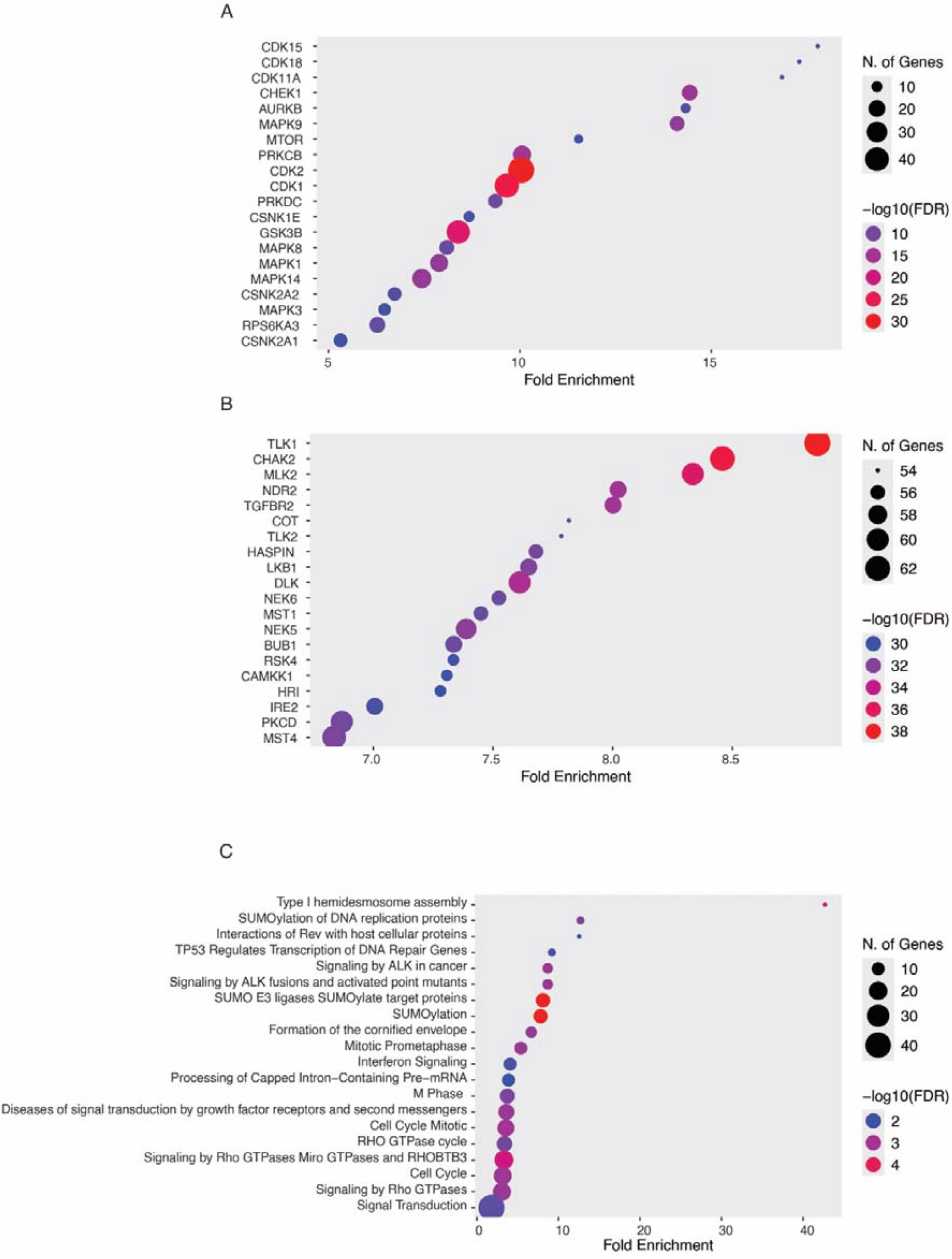
Enrichment analysis reveals coordinated suppression of cell cycle and checkpoint kinase networks following AME treatment. (A, B) Kinase enrichment analysis of hypophosphorylated phosphoproteins performed using ShinyGO 0.85.1 with the Kinase Library 2023 and KEA 2015 databases. Substrates associated with multiple cyclin-dependent kinases (CDK1, CDK2, CDK7, CDK11A, CDK15, and CDK18), as well as mitotic and checkpoint regulators including AURKB, CHEK1, CHEK2, and PRKDC, were significantly enriched. In addition, hypophosphorylated substrates corresponding to mitogen-activated protein kinases (MAPK1, MAPK3, MAPK8, MAPK9, and MAPK14) and mTOR were prominently overrepresented, indicating broad attenuation of proliferative and survival-associated kinase signaling. (C) Reactome pathway enrichment analysis of hypophosphorylated proteins showing significant association with cell cycle–related pathways, including mitotic progression, mitotic prometaphase, and M-phase regulation, as well as TP53-mediated DNA damage response, SUMOylation of DNA replication proteins, and DNA damage checkpoint signaling. Enrichment analyses were performed using ShinyGO 0.85.1, and statistically significant terms were identified based on adjusted p-values as described in the Methods section.

Pathway enrichment analysis using the Reactome database in ShinyGO 0.85.1 online tool further demonstrated that hypophosphorylated proteins were significantly associated with cell cycle related pathways, including cell cycle mitotic progression, mitotic prometaphase, and M-phase regulation. Notably, pathways involved in TP53-mediated regulation of DNA repair, SUMOylation of DNA replication proteins, and DNA damage checkpoint signaling were also enriched, suggesting activation of growth-suppressive and checkpoint-associated regulatory programs following AME treatment (Fig. 9C).

Collectively, these analyses reveal that AME treatment induces a coordinated hypophosphorylation of kinase networks governing cell cycle progression, mitotic control, and checkpoint signaling, providing a phosphoproteomic basis for the observed anti-proliferative effects of AME in OSCC cells.

## Discussion

Previous in vitro and in vivo studies have demonstrated that *Annona muricata* extracts exert anticancer and antitumor activities across multiple cancer types, including breast, liver, prostate, pancreatic, lung, and colon cancers [9, 25-27]. The ability of *A. muricata* extracts to induce selective cytotoxicity toward cancer cells has generated considerable interest in their potential therapeutic applications [9, 28, 29]. Notably, extracts from *A. muricata* leaves have been widely used in traditional medicine and are often referreds to as a “cancer killer.” Consistent with these observations, Najmuddin *et al*. reported that *A. muricata* leaf extracts exhibited cytotoxic effects in breast cancer cell lines while exerting minimal toxicity toward normal breast epithelial cells, highlighting their potential cancer selectivity [29].

In the present study, we investigated the anticancer effects of methanolic *A. muricata* extract (AME) in CAL-27 cells, a widely used oral squamous cell carcinoma (OSCC) model. We observed that AME treatment significantly reduced cell proliferation, migration, and clonogenic survival, indicating potent anti-tumor activity. Among the tested extracts, the methanolic fraction exhibited greater cytotoxicity than the aqueous extract, consistent with the enrichment of bioactive phytochemicals in organic solvents.

Phosphorylation is a critical post-translational modification that regulates diverse cellular processes under physiological conditions, whereas dysregulated phosphorylation signaling is a hallmark of cancer [30-32]. Aberrant kinase activity frequently drives uncontrolled proliferation and survival, making phosphorylation-dependent pathways attractive targets for anticancer therapy [33, 34]. Although previous studies have suggested that *A. muricata* extracts can modulate oncogenic signaling, the underlying molecular mechanisms have remained poorly defined. To address this gap, we performed a quantitative phosphoproteomic analysis to systematically characterize signaling alterations induced by AME treatment. Our data revealed that the majority of phosphoproteomic changes occurred at later time points (1–2 h), suggesting a coordinated rewiring of intracellular signaling rather than an immediate stress response. Pathway enrichment analysis demonstrated that hypophosphorylated proteins were predominantly associated with cell cycle regulation, consistent with the observed suppression of proliferation.

Cell cycle progression is tightly controlled by cyclin-dependent kinases (CDKs), and dysregulated CDK activity is a well-established driver of tumorigenesis. Both hyperactivation and inappropriate suppression of CDKs can disrupt cell cycle fidelity and contribute to malignant transformation [35-38]. In this study, AME treatment resulted in significant hypophosphorylation of CDK1, CDK2, and CDK5 substrates, indicating attenuation of CDK-dependent signaling. These findings suggest that AME interferes with cell cycle progression, thereby suppressing OSCC cell growth.

In addition to CDKs, components of the mitogen-activated protein kinase (MAPK) pathway were prominently hypophosphorylated following AME treatment. The MAPK signaling cascade plays a central role in oncogenesis, tumor progression, and therapeutic resistance, and aberrant MAPK activation has been reported in multiple cancers, including OSCC[39-41]. We observed reduced phosphorylation of MAPK1, MAPK10, MAPK14, MAP2K1, and MAP2K6, consistent with inhibition of proliferative and survival-associated signaling. These phosphoproteomic findings align with our biochemical data demonstrating reduced ERK and AKT phosphorylation, further supporting the suppression of oncogenic signaling pathways.

Among the consistently hypophosphorylated signaling proteins identified across multiple time points were EGFR, BAIAP2L1, MARCKS, PKP2, ARAF, ARHGEF16, MAPRE1, CDK12, and NUCKS1. EGFR is a receptor tyrosine kinase that orchestrates key signaling pathways governing proliferation, survival, and differentiation [42, 43]. Previous studies have shown that *A. muricata* derived compounds can interact with EGFR signaling to induce cell cycle arrest and apoptosis while inhibiting migration and metastasis [44]. In agreement with these reports, we observed reduced EGFR phosphorylation following AME treatment, suggesting attenuation of EGFR-driven signaling in OSCC cells.

Several EGFR-associated regulatory proteins were also hypophosphorylated. PKP2 has been reported to promote EGFR dimerization and downstream signaling, thereby facilitating tumor progression [45]. Similarly, BAIAP2L1 functions as an adaptor protein that enhances EGFR-mediated activation of ERK and PI3K pathways [46]. Hypophosphorylation of these proteins upon AME treatment suggests disruption of EGFR-centered signaling complexes, leading to reduced downstream oncogenic signaling.

MARCKS is another critical signaling regulator whose phosphorylation status has been linked to PI3K/AKT pathway activation, tumor progression, and therapeutic resistance [47]. Elevated MARCKS phosphorylation has been associated with poor prognosis in multiple cancers, including OSCC and lung cancer, whereas inhibition of MARCKS phosphorylation enhances sensitivity to chemotherapy and radiotherapy [48, 49]. In this study, AME-induced hypophosphorylation of MARCKS suggests a potential mechanism by which AME suppresses PI3K/AKT signaling and enhances growth inhibition in OSCC cells.

To further contextualize the global signaling impact of AME, we performed integrative kinase and pathway enrichment analyses focused specifically on hypophosphorylated phosphoproteins. These analyses revealed a striking overrepresentation of substrates associated with multiple cyclin-dependent kinases (CDK1, CDK2, CDK7, CDK11A, CDK15, and CDK18), together with key mitotic and checkpoint regulators including AURKB, CHEK1, CHEK2, and PRKDC. These kinases constitute the core machinery that governs cell cycle progression, mitotic fidelity, and DNA damage surveillance, and their dysregulation is a hallmark of malignant transformation and therapeutic resistance [32, 34, 36, 37]. The coordinated hypophosphorylation of their downstream substrates following AME treatment suggests a broad suppression of proliferative and checkpoint-adaptive signaling programs that are essential for OSCC cell survival and division.

In parallel, enrichment of hypophosphorylated substrates linked to MAPK family members (MAPK1, MAPK3, MAPK8, MAPK9, and MAPK14) and mTOR further supports the notion that AME exerts multi-layered inhibition of oncogenic signaling cascades that converge on cell growth, metabolism, and survival. Aberrant MAPK and mTOR activation is frequently observed in OSCC and has been implicated in aggressive tumor behavior, metastatic progression, and resistance to chemotherapy and radiotherapy [39-41]. The observed attenuation of these kinase networks is therefore consistent with the pronounced anti-proliferative and anti-migratory effects of AME in CAL-27 cells.

Pathway-level enrichment using the Reactome database further demonstrated that hypophosphorylated proteins were predominantly associated with mitotic progression, prometaphase regulation, and M-phase control, underscoring a profound disruption of cell cycle execution following AME treatment. In addition, enrichment of TP53-mediated DNA damage response pathways, SUMOylation of DNA replication proteins, and DNA damage checkpoint signaling suggests that AME not only suppresses proliferative signaling but may also reinforce growth-suppressive and genome-protective programs. Together, these findings indicate that AME induces a coordinated collapse of oncogenic kinase networks while simultaneously engaging checkpoint-associated regulatory circuits, thereby creating a cellular environment incompatible with sustained tumor cell proliferation.

Collectively, our findings demonstrate that AME induces a coordinated and systems-level rewiring of oncogenic signaling networks in OSCC cells through widespread hypophosphorylation of key regulatory proteins. Rather than acting on a single molecular target, AME suppresses multiple interconnected kinase pathways governing cell cycle progression, mitotic control, DNA damage checkpoint regulation, and EGFR/MAPK and PI3K/AKT signaling. The concerted inhibition of CDK- and MAPK-driven proliferative programs, together with attenuation of EGFR-associated signaling complexes and suppression of MARCKS-dependent PI3K/AKT activation, provides a mechanistic basis for the observed anti-proliferative, anti-migratory, and clonogenic inhibitory effects of AME. These data suggest that AME enforces a growth-restrictive cellular state by collapsing oncogenic kinase circuitry while reinforcing checkpoint-associated regulatory programs. Our study therefore establishes a phosphoproteomic framework for understanding the anticancer activity of *Annona muricata* and highlights the therapeutic potential of phytochemical-mediated modulation of phosphorylation-dependent signaling networks in OSCC.

## Conclusions

Monitoring phosphorylation changes at the proteome level following *Annona muricata* treatment enabled a systematic investigation of key signaling pathways involved in oral squamous cell carcinoma. This study demonstrates that *A. muricata* exhibits significant anticancer potential in OSCC by modulating phosphorylation-dependent signaling networks that govern cell proliferation, survival, migration, and differentiation. The observed alterations in kinase-associated signaling pathways suggest that AME induces broad rewiring of oncogenic signaling rather than acting through a single molecular target.

These findings highlight several dysregulated kinases and signaling regulators as potential candidates for therapeutic intervention in OSCC and provide a mechanistic framework for understanding the anticancer effects of *A. muricata*. However, further studies are required to evaluate crude extracts and isolated bioactive compounds to determine their potency, optimal dosage, precise mechanisms of action, long-term safety, and possible adverse effects. Such investigations will be essential for translating the anticancer potential of *A. muricata* into clinically relevant therapeutic strategies.

## Supporting information

Supplementary Figure 1. Gene Ontology (GO) analysis of differentially regulated proteins upon AME treatment in CAL-27 cells to classify the proteins b

Supplementary Table 1. List of phosphopeptides identified and quantified in TMT based quantitative phosphoproteomic approach in CAL-27 cells upon AME

Supplementary Table 2. List of differentially phosphorylated peptides upon AME treatment in CAL-27 cells.

## Abbreviations

BCA: bicinchoninic acid
CDK: cyclin dependent kinase
CSNK1A1: casein kinase 1 alpha 1
DPPH: 2,2-diphenylpicrylhydrazyl
EGFR: epidermal growth factor receptor
FBS: fetal bovine serum
LC-MS/MS: liquid chromatography-mass spectrometry/mass spectrometry
MAPK: mitogen-activated protein kinase
MTOR: mammalian target of rapamycin
MTT: 3-(4,5-dimethylthiazol-2-yl)-2,5-diphenyl tetrazolium bromide
PKC: protein kinase C
PRKCD: protein kinase C delta
ROCK1: Rho associated coiled-coil containing protein kinase 1
TEABC: triethyl ammonium benzyl chloride.

## Data availability statement

The MS raw data and the Proteome Discoverer-searched data were submitted to the ProteomeXchange Consortium (http://proteomecentral.proteomexchange.org) via the PRIDE repository with the data set identifier PXD033096.

## CRediT author statement

**Shobha Dagamajalu**: Writing original draft, Investigation, Supervision, Methodology. **Mohd Altaf Najar**: Writing original draft, Methodology, Visualization, Data analysis. **Devasahayam Arokia Balaya Rex**: Data analysis, Visualization. **Prashant Kumar Modi**:

Methodology. **Suchitha G. P**: Methodology. **Amrutha S**: Methodology. **T. S. Keshava Prasad**: Resources, Project administration, Supervision.

## Acknowledgments

The authors gratefully acknowledge Yenepoya (Deemed to be University) for providing the infrastructure and state-of-the-art mass spectrometry facility required to conduct this study exclusively within our institution. We also acknowledge the support provided by the Department of Biotechnology (DBT) through the National Facility grant under the project “Skill Development in Mass Spectrometry-based Metabolomics Technology BIC” (BT/PR40202/BTIS/137/53/2023).

