## Supplementary Figure 1. Gene Ontology (GO) analysis of differentially regulated proteins upon AME treatment in CAL-27 cells to classify the proteins b for "Phosphoproteomic profiling reveals signaling pathways modulated by *Annona muricata* leaf extract in oral adenosquamous carcinoma cells"

Shobha Dagamajalu. Ph.D.,

Associate Professor

Center for Systems Biology and Molecular Medicine

Yenepoya Research Centre

Yenepoya (Deemed to be University)

Mangalore 575018, India

T. S. Keshava Prasad. Ph.D.,

Professor and Deputy Director

Center for Systems Biology and Molecular Medicine

Yenepoya Research Centre

Yenepoya (Deemed to be University)

Mangalore 575018, India

**ORCID**

Mohd Altaf Najar	0000-0002-3044-3772

Devasahayam Arokia Balaya Rex	0000-0002-9556-3150

Prashant Kumar Modi	0000-0002-4817-3379

Suchitha G. P	0000-0002-4930-2590

Amrutha S	0000-0001-7696-0903

T. S. Keshava Prasad	0000-0002-6206-2384

Shobha Dagamajalu	0000-0002-0899-2839

**Institutional Email IDs**

Mohd Altaf Najar

Devasahayam Arokia Balaya Rex

Prashant Kumar Modi

Suchitha G. P

Amrutha, S

T. S. Keshava Prasad

Shobha Dagamajalu

**Abstract**

Phosphorylation driven dysregulation of intracellular signaling networks is a central feature of cancer initiation, progression, and therapeutic resistance. Although *Annona muricata* leaf extracts have demonstrated anticancer activity across multiple experimental models, the underlying molecular mechanisms particularly at the level of phosphorylation dependent signaling remain poorly understood. In this study, we employed a tandem mass tag TMT-based quantitative phosphoproteomic approach to systematically characterize signaling alterations induced by methanolic *Annona muricata*leaf extract (AME) in oral squamous cell carcinoma (OSCC) CAL-27 cells.

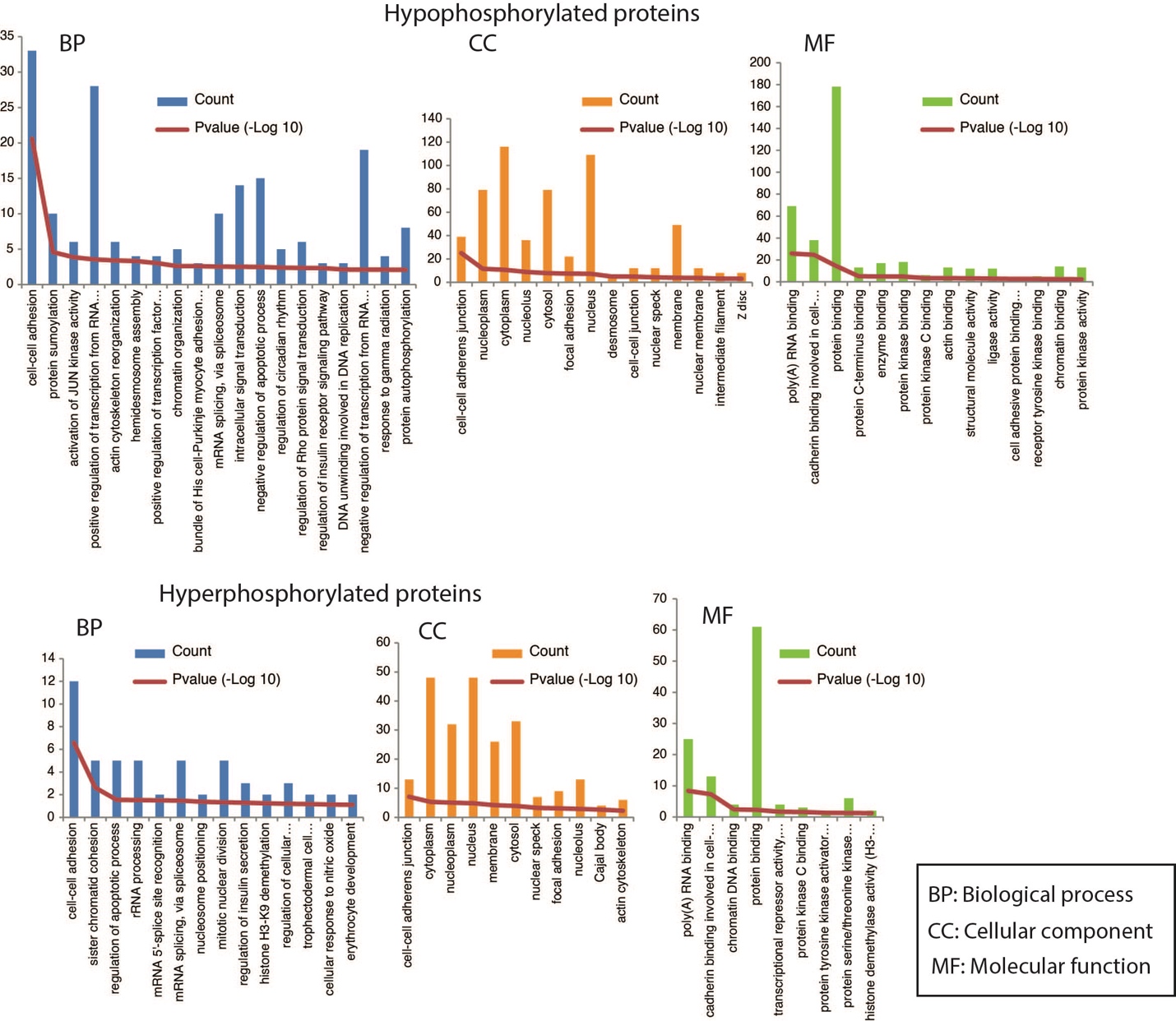

**Supplementary Figure 1.** Gene Ontology (GO) analysis of differentially regulated proteins upon AME treatment in CAL-27 cells to classify the proteins based on biological process, cellular component and molecular function.

**Supplementary Table 1.** List of phosphopeptides identified and quantified in TMT based quantitative phosphoproteomic approach in CAL-27 cells upon AME treatment.

**Supplementary Table 2.** List of differentially phosphorylated peptides upon AME treatment in CAL-27 cells.
